# CXXC1 is redundant for normal DNA double-strand break formation and meiotic recombination in mouse

**DOI:** 10.1101/305508

**Authors:** Hui Tian, Timothy Billings, Petko M. Petkov

## Abstract

In most mammals, including mice and humans, meiotic recombination is determined by the meiosis specific histone methytransferase PRDM9, which binds to specific DNA sequences and trimethylates histone 3 at lysine-4 and lysine-36 at the adjacent nucleosomes. These actions ensure successful DNA double strand break initiation and repair that occur on the proteinaceous structure forming the chromosome axis. The process of hotspot association with the axis after their activation by PRDM9 is poorly understood. Previously, we and others have identified CXXC1, an ortholog of *S. cerevisiae* Spp1 in mammals, as a PRDM9 interactor. In yeast, Spp1 is a histone methyl reader that links H3K4me3 sites with the recombination machinery, promoting DSB formation. Here we investigated whether CXXC1 has a similar function in mouse meiosis. We found that CXXC1 is co-expressed and interacts with PRDM9 in mouse spermatocytes. To investigate the meiotic function of CXXC1, we created a *Cxxc1* conditional knockout mouse to deplete CXXC1 before the onset of meiosis. Surprisingly, knockout mice were fertile, and the loss of CXXC1 in spermatocytes had no effect on hotspot trimethylation activity, double-strand break formation or repair. Our results demonstrate that CXXC1 is not an essential link between recombination hotspot sites and DSB machinery and that the hotspot recognition pathway in mouse is independent of CXXC1.

**Author Summary:** Meiotic recombination increases genetic diversity by ensuring novel combination of alleles passing onto the next generation correctly. In most mammals, the meiotic recombination sites are determined by histone methyltransferase PRDM9. These sites subsequently become associated with the chromosome axis with the participation of additional proteins and undergo double strand breaks, which are repaired by homologous recombination. In *Saccharomyces cerevisiae*, Spp1 (ortholog of CXXC1) binds to methylated H3K4 and connects these sites with chromosome axis promoting DSB formation. However, our data suggest that even though CXXC1 interacts with PRDM9 in male germ cells, it does not play a crucial role in mouse meiotic recombination. These results indicate that, unlike in *S. cerevisiae*, a recombination initiation pathway that includes CXXC1 could only serve as a non-essential pathway in mouse meiotic recombination.

## Introduction

Meiotic recombination ensures production of fertile gametes with a correct haploid chromosome number and genetic diversity [1]. In most of mammals, the meiotic recombination sites are restricted to 1-2 kb regions, locations of which are determined by the DNA binding histone methyltransferase PRDM9 [2–4]. Recombination initiates when PRDM9 binds to hotspots sequences with its zinc finger domain and trimethylates histone 3 at lysine 4 (H3K4me3) and lysine 36 (H3K36me3) resulting in formation of a nucleosome-depleted region [2–6]. DNA double strand breaks (DSB) are created at the nucleosome-depleted regions of activated hotspots [7–10], and eventually repaired as either crossovers or non-crossover conversions. Cytological staining for several proteins associated with DSB processing in early meiotic prophase show that, from the earliest time of their detection, DSB are associated with a proteinaceous structure known as chromosome axis [11–14]. We have previously shown that PRMD9 is associated, but not directly interacting, with chromosome axis elements such as phosphorylated REC8 (pREC8) and SYCP3 in spermatocytes [15]. Efficient trimethylation at hotspots and correct association between PRDM9 and certain chromosomal axis elements are crucial for normal DSB formation and repair [16]. However, we currently do not have detailed knowledge of the proteins and molecular mechanisms participating in hotspot association with chromosome axis.

In *Saccharomyces cerevisiae*, which has no PRDM9, the PHD zinc finger protein Spp1, a member of COMPASS (Complex associated with Set1), acts as a histone H3K4 methyl reader and promotes meiotic DSB formation at the existing H3K4me3 sites, such as promoters [17]. Spp1 is predominantly located on the chromosome axes and connects H3K4 trimethylated sites with the axis protein Mer2 to stimulate Spo11 dependent DSB formation [17,18]. Recent study showed that Spp1 function in tethering DSB sites to chromosome axes and ensuring efficient DSB formation is independent of its function as a COMPASS complex member [19].

CxxC finger protein 1 (CXXC1, CFP1, CGBP) is an ortholog of *S. cerevisiae* Spp1 in mammals. In somatic cells, CXXC1 binds to both unmethylated CpGs and SETD1, which is required for trimethylation of H3K4 at CpG islands [20]. CXXC1 is crucial for embryonic stem cell maintaining and development [21,22]. Knockout of *Cxxc1* in mouse results in lethality at the early embryonic stages [23]. We have reported that CXXC1 interacts with PRDM9 *in vitro* [15]. This interaction has recently been confirmed by another group, which also reported that CXXC1 interacts with the chromosome axis element IHO1 by yeast two-hybrid assay [24]. IHO1 is considered to be the ortholog of yeast Mer2 and is known to be essential for ensuring efficient DSB formation [25], therefore it is possible that CXXC1-IHO1 interaction serves the same function in mammalian meiosis as their orthologs Spp1-Mer2 in yeast. However, the function of CXXC1 in mammalian meiosis has not been characterized so far. It has been unclear whether CXXC1 binds to PRDM9 in germ cells and whether it participates in meiotic recombination initiation in any way, either as a partner of PRDM9 or as a methyl reader of H3K4me3/H3K36me3 marks that PRDM9 imposes at the nucleosomes surrounding the recombination hotspots.

To address whether and how CXXC1 functions in meiotic recombination, we created a *Cxxc1* conditional knockout mouse model and deleted *Cxxc1* at the onset of meiosis. We found that CXXC1 is co-expressed with PRDM9 and indeed interacts with it in spermatocytes. However, loss of CXXC1 does not affect normal meiotic recombination process. Our study demonstrates that the presence of CXXC1 in mouse meiosis is redundant, and unlike its *S. cerevisiae* ortholog Spp1, CXXC1 does not appear to be a key factor for the DSB formation.

## Results

### CXXC1 interacts with PRDM9 in spermatocytes

We tested whether CXXC1 interacts with PRDM9 *in vivo* by co-immunoprecipitation (co-IP) from spermatocytes isolated from 14 dpp B6 testis using antibody against PRDM9. We found that CXXC1 indeed interacts with PRDM9 in spermatocytes (Fig 1A). However, the interaction was not as strong as PRDM9’s predominant interactor EWSR1 (Fig 1A) [15], which raised the possibility that the interaction between CXXC1 and PRDM9 could be mediated by the stronger PRDM9 interactors. To test whether this is the case, we performed co-IP with EWSR1 and did not detect any interaction with CXXC1 in testicular extract (Fig 1B). To further test the interactions between the three proteins, CXXC1, PRDM9 and EWSR1, we co-expressed Myc-tagged mouse CXXC1, HA-tagged EWSR1 and Flag-tagged PRDM9 proteins in human embryonal kidney 293 (HEK293) cells, and performed co-IP with antibodies against HA or Myc tags. Both EWSR1 and CXXC1 immunoprecipitated PRDM9 under these conditions, but there was no interaction between CXXC1 and EWSR1 in the presence or absence of PRDM9 (Fig 1C).

**Fig 1.**
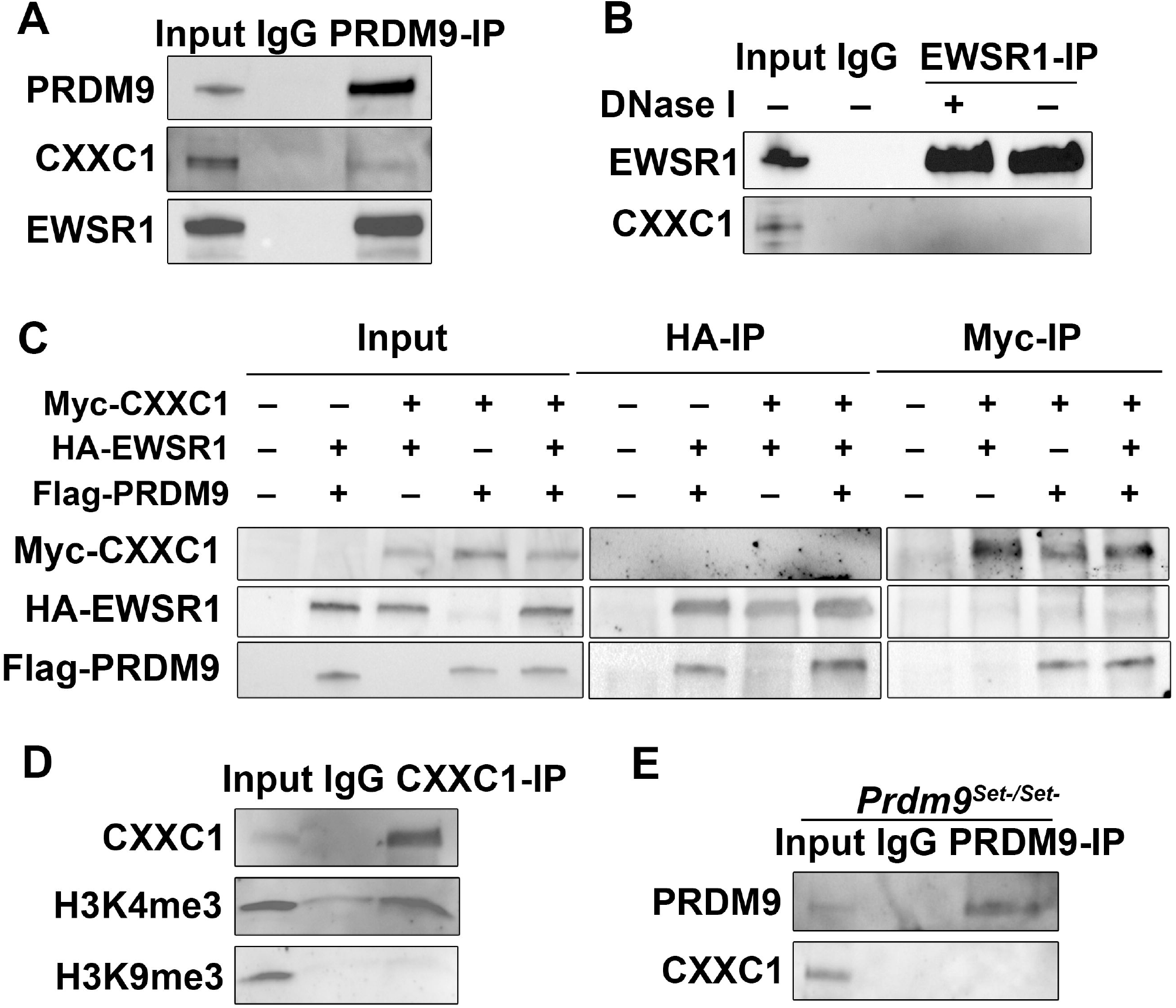
CXXC1 interacts with PRDM9 in spermatocytes. (A) Co-IP with PRDM9 from 14 dpp B6 testicular extract. Staining for PRDM9, CXXC1, EWSR1 and CTCF. In each blot, lane 1- input; lane 2 – co-IP with non-immune IgG; lane 3 – co-IP with antibody against PRDM9. (B) Co-IP with EWSR1 from 14 dpp B6 testicular extract. Staining for CXXC1 and EWSR1. In each blot, lane 1- input; lane 2 – co-IP with non-immune IgG; lane 3 – co-IP with anti-EWSR1 after DNase I treatment; co-IP with anti-EWSR1 without DNase I treatment. (C) Myc tagged CXXC1, HA tagged EWSR1 and Flag tagged PRDM9 were transfected into HEK293 cells. Co-IP with HA or Myc antibody was performed. The indicated proteins are stained. (D) Co-IP with CXXC1 from 14 dpp B6 testicular extract showed there is interaction between CXXC1 and H3K4me3. H3K9me3 is used as a negative control. (E) Co-IP with PRDM9 from 14 dpp *Prdm9^Set-/Set-^* testicular extract. Staining for PRDM9 and CXXC1. In each blot, lane 1- input; lane 2 – co-IP with non-immune IgG; lane 3 – co-IP with antibody against PRDM9.

Several reports have shown that Spp1, the yeast ortholog of CXXC1, binds to H3K4me3 and tethers H3K4-trimethylated recombination hotspots to the chromosome axis [17–19]. To test whether CXXC1 binds to H3K4me3 in mouse spermatocytes, we performed CXXC1 co-IP from B6 testicular extract. Indeed, we detected CXXC1 interaction with H3K4me3 but not with the closed chromatin marker H3K9me3 (Fig 1D). These data raise the possibility that the *in vivo* interaction between PRDM9 and CXXC1 could be mediated by trimethylated H3K4. To further investigate that, we performed PRDM9 co-IP from the testicular extract isolated from a PRDM9 PR/SET domain mutation mouse model (*Prdm9^Set-/Set-^*), in which hotspots are not trimethylated (unpublished data). We did not detect any interaction between CXXC1 and PRDM9 in the mutant testicular extract (Fig 1E), which indicates that CXXC1 binding to PRDM9 may be mediated or at least facilitated by trimethylated H3K4 at hotspots.

These results indicate that CXXC1 and EWSR1 form separate complexes with PRDM9. They also indicate that although CXXC1 interacts with PRDM9 *in vivo*, it is not a predominant interactor of PRDM9, and that their interaction could be mediated by other proteins such as histone 3 trimethylated at lysine-4.

### CXXC1 is co-expressed with PRDM9 in leptonema and zygonema

In seminiferous tubules of mouse testis, CXXC1 is expressed in both germ cells and Sertoli cells (Fig 2A, top panel). CXXC1 showed high expression in spermatogonia, low expression in leptonema and zygonema, and then again high expression in pachynema and diplonema, decreasing to undetectable levels in spermatids (Fig 2A, top panel).

**Fig 2.**
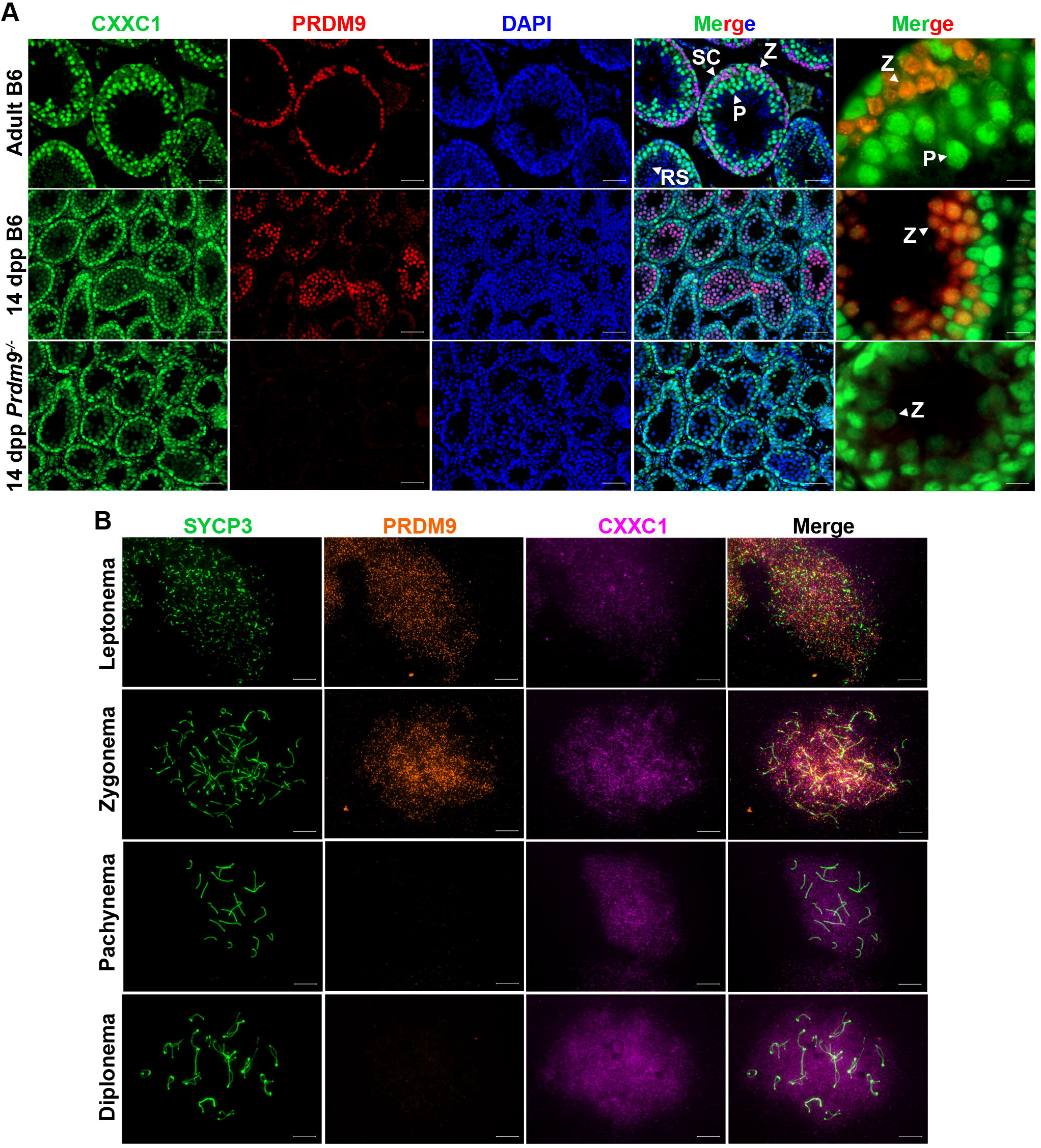
CXXC1 is co-expressed in the spermatocytes with PRDM9. (A) Co-immunofluorescence staining of CXXC1 and PRDM9 on adult B6, 14-dpp B6 and 14-dpp *Prdm9^Set-/Set-^* seminiferous tubule cross sections. Green, CXXC1; red, PRDM9; blue, DAPI. SC, Sertoli cell; Z, zygonema; P, pachynema; RS, round spermatid. Scale bars, first 4 columns: 50 μm, last column: 10 μm. (B) Immunostaining of CXXC1 and PRDM9 on chromosome spreads from adult B6. Green, SYCP3; orange, PRDM9; magenta, CXXC1. Scale bar, 10 μm.

Previous reports showed that PRDM9 is present only in leptonema and zygonema during meiosis [26]. Double staining for PRDM9 and CXXC1 showed co-expression of these two proteins in nuclei from stage X seminiferous tubules (Fig 2A, top panel) and in 14 dpp testis (Fig 2A, middle panel), when the majority of spermatocytes are at leptotene and zygotene stages. Since CXXC1 interacts with PRDM9 *in vivo* (Fig 1A), we performed CXXC1 staining in *Prdm9* knockout mouse testis (*Prdm9^-/-^*) to determine whether CXXC1 localization could be affected by the absence of PRDM9. In this mutant, CXXC1 showed the same localization pattern as in controls (Fig 2A, bottom panel). The co-expression pattern of PRDM9 and CXXC1 in leptonema and zygonema was further confirmed by chromosome spreads, where CXXC1 showed diffused signal over the entire nuclear region (Fig 2B).

These results suggest that CXXC1 is co-expressed with PRDM9 in leptonema and zygonema, and that its expression and localization are not affected by the presence or absence of PRDM9.

### Male *Cxxc1* knockout mice are fertile

To test whether CXXC1 is involved in spermatogenesis, we generated a conditional knockout (CKO) model using CRISPR/Cas9 to insert loxP sites flanking exon 2 and 3 of *Cxxc1* in C57BL/6J. We bred early spermatogonia-specific knockout mice (*Cxxc1^loxp/Δ;Stra8-iCre^*, hereafter *Cxxc1* CKO) by crossing the *Cxxc1^loxp/loxP^* mice with *Stra8-iCre* mice. Western blot confirmed that in knockout testes, the protein level of CXXC1 is reduced (Fig 3A). CXXC1 was absent in spermatocytes of the CKO, but present in spermatogonia and Sertoli cells (Fig. 3B).

**Fig 3.**
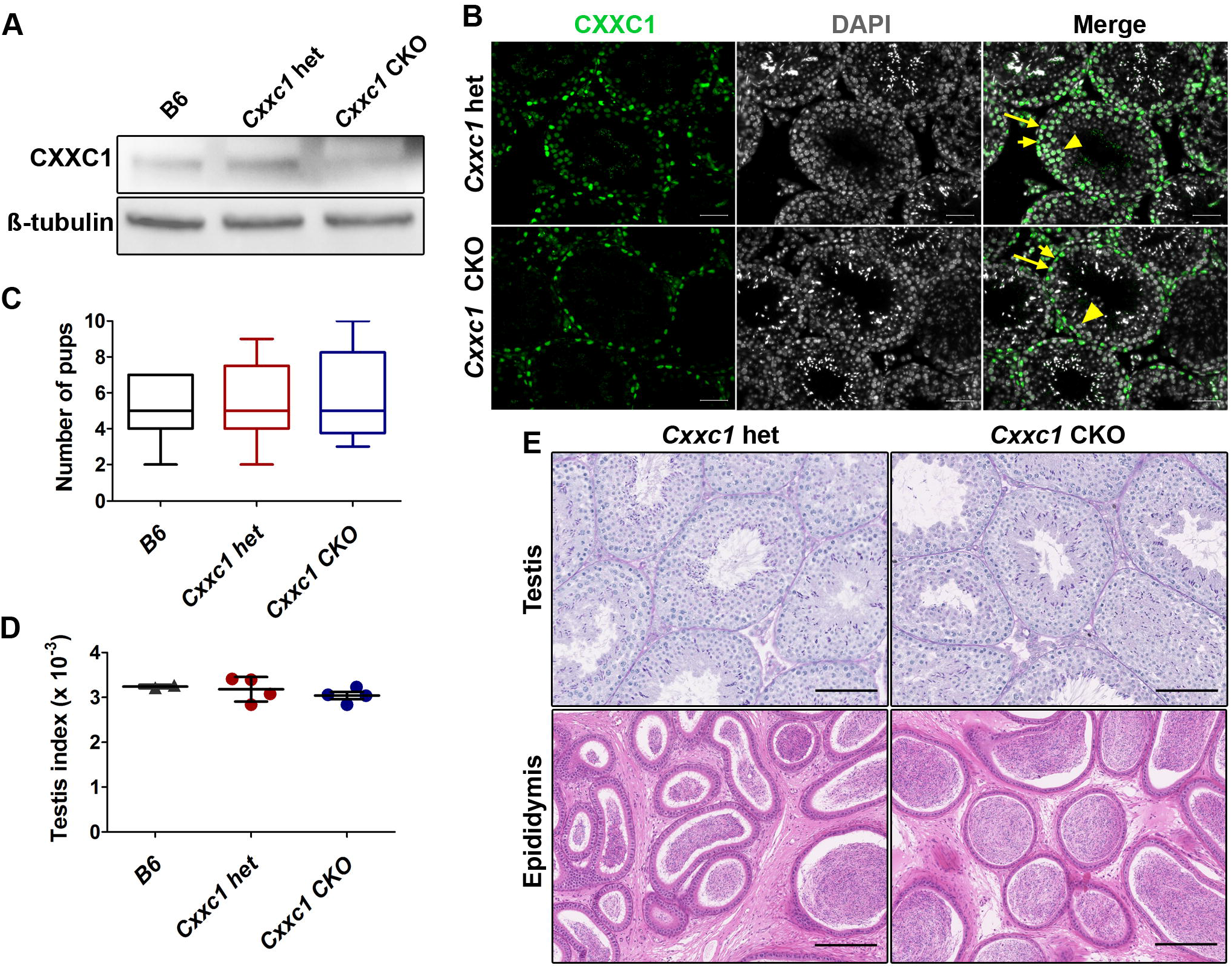
Knockout of CXXC1 does not affect male fertility or testis histology. (A) Western blot of CXXC1 with adult B6, *Cxxc1* het and CKO testicular extract. ß-tubulin was used as internal loading control. (B) Immunostaining of CXXC1 on *Cxxc1* het and CKO seminiferous tubule cross sections. Scale bar, 50 μm. Long arrows, Sertoli cells; short arrows, spermatogonia; arrowhead, spermatocytes. (C) Fertility tests in B6, *Cxxc1* het and CKO mice. The number of viable pups was shown. (D) Testis index (testis weight/body weight) in B6, *Cxxc1* het and CKO mice. (E) Histology of testis and epididymis in *Cxxc1* control and CKO mice. Top panels, PAS staining of seminiferous tubule sections; scale bar, 100 μm. Bottom panels, H&E staining of epididymis sections; scale bar, 200 μm. Left panels, het control; right panels, *Cxxc1* CKO.

We performed fertility test with two CKO mice. To our surprise, both CKO mice were fertile and produced similar number of viable progeny compared to the heterozygous (het) and B6 controls (Fig 3C). Testis index (testis weight/body weight) was the same in CKO as in het and wild type B6 controls (Fig. 3D). Histology of testis and epididymis from CKO mice showed no detectable spermatogenesis defects (Fig 3E). In contrast, *Cxxc1* oocyte-specific knockout female mice with *Ddx4*-Cre are sterile – no viable pup produced from homozygous knockout (*Cxxc1^loxp/Δ;Ddx4-Cre^*) mating test, while the heterozygous control (*Cxxc1^loxp/+;Ddx4-Cre^*) mating produced normal number of pups (5.3 ± 1.7); however, their sterility is due to early embryonic developmental deficiency and not meiotic defects [27].

### Expression and function of PRDM9 remains normal in *Cxxc1* CKO

We further tested whether CXXC1 affects the localization, the expression pattern, or the function of PRDM9. Localization of PRDM9 in seminiferous tubules was preserved in CKO as determined by immunostaining (Fig 4A). In addition, the expression pattern of PRDM9 in leptonema and zygonema was not affected in the absence of CXXC1 (Fig 4B).

**Fig 4.**
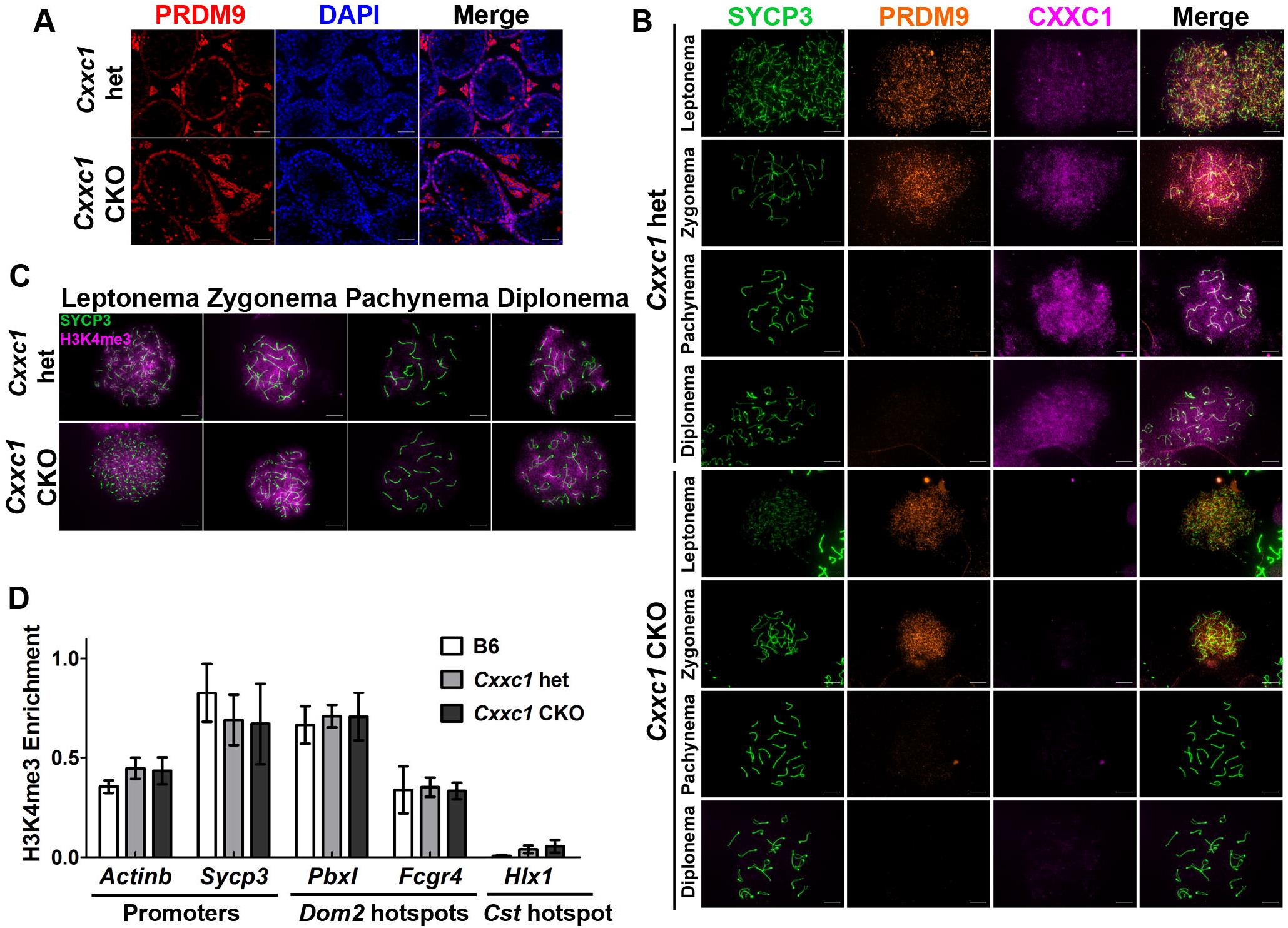
PRDM9 expression and catalytic function are not impaired in *Cxxc1* CKO. (A) Immunostaining of PRDM9 in adult *Cxxc1* het and CKO seminiferous tubules. Scale bar, 50 μm. (B) Co-immunostaining of CXXC1 and PRDM9 on chromosome spreads from adult *Cxxc1* het and CKO mice. Green, SYCP3; orange, PRDM9; magenta, CXXC1. First 4 rows, het control; last 4 rowls, *Cxxc1* CKO. Scale bar, 10 μm. (C) Immunostaining of H3K4me3 in adult *Cxxc1* het and CKO seminiferous tubules. Green, SYCP3; magenta, H3K4me3. Scale bar, 10 μm. (D) H3K4me3 ChIP-qPCR with chromatin isolated form *Cxxc1* het and CKO mice. Promoter regions form *Actinb* and *Sycp3, Dom2* hotspots *Pbxl* and *Fcgr4* were amplified. *Cst* hotspot *Hlxl* was used as a negative control. Bars present mean ± SD of three biological replicates.

To test whether lack of CXXC1 affects PRDM9 methyltransferase function, we first compared H3K4me3 patterns in control and *Cxxc1* CKO mice. Both control and CKO chromosome spreads showed abundant H3K4me3 signal in leptonema and zygonema, lower signal in pachynema and increased signal in diplonema (Fig 4C). In addition, the H3K4me3 staining on cross sections of CKO testis showed no decrease (Fig S1). These data indicate that the hotspot trimethylation and transcriptional activation in spermatocytes are not affected by the loss of CXXC1.

Second, we determined whether loss of CXXC1 affects PRDM9 binding and its methytransferase activity at individual hotspots by H3K4me3 ChIP-qPCR. We found that H3K4me3 enrichment at *Dom2* hotspots *PbxI* and *Fcgr4* was not different in B6 control, *Cxxc1* heterozygous and CKO. We measured as a control the H3K4me3 enrichment at promoter regions of housekeeping gene *Actinb* and meiotic specific gene *Sycp3*, which are not PRDM9-dependent. These were not changed as well (Fig 4D).

These results suggest that loss of CXXC1 does not affect PRDM9 expression, binding to hotspots, or its catalytic function. Therefore, CXXC1 is not required for PRDM9-dependent hotspot activation.

### Meiotic DSB formation and repair are normal in *Cxxc1* CKO

Next, we tested whether lack of CXXC1 affects chromosome synapsis, DSB formation and repair process. Co-staining of SYCP1 and SYCP3 showed normal synapsis in all autosomes at pachytene stage in CKO spermatocytes. We did not detect increased chromosome asynapsis in the CKO spermatocytes compared to controls (Fig 5A).

**Fig 5.**
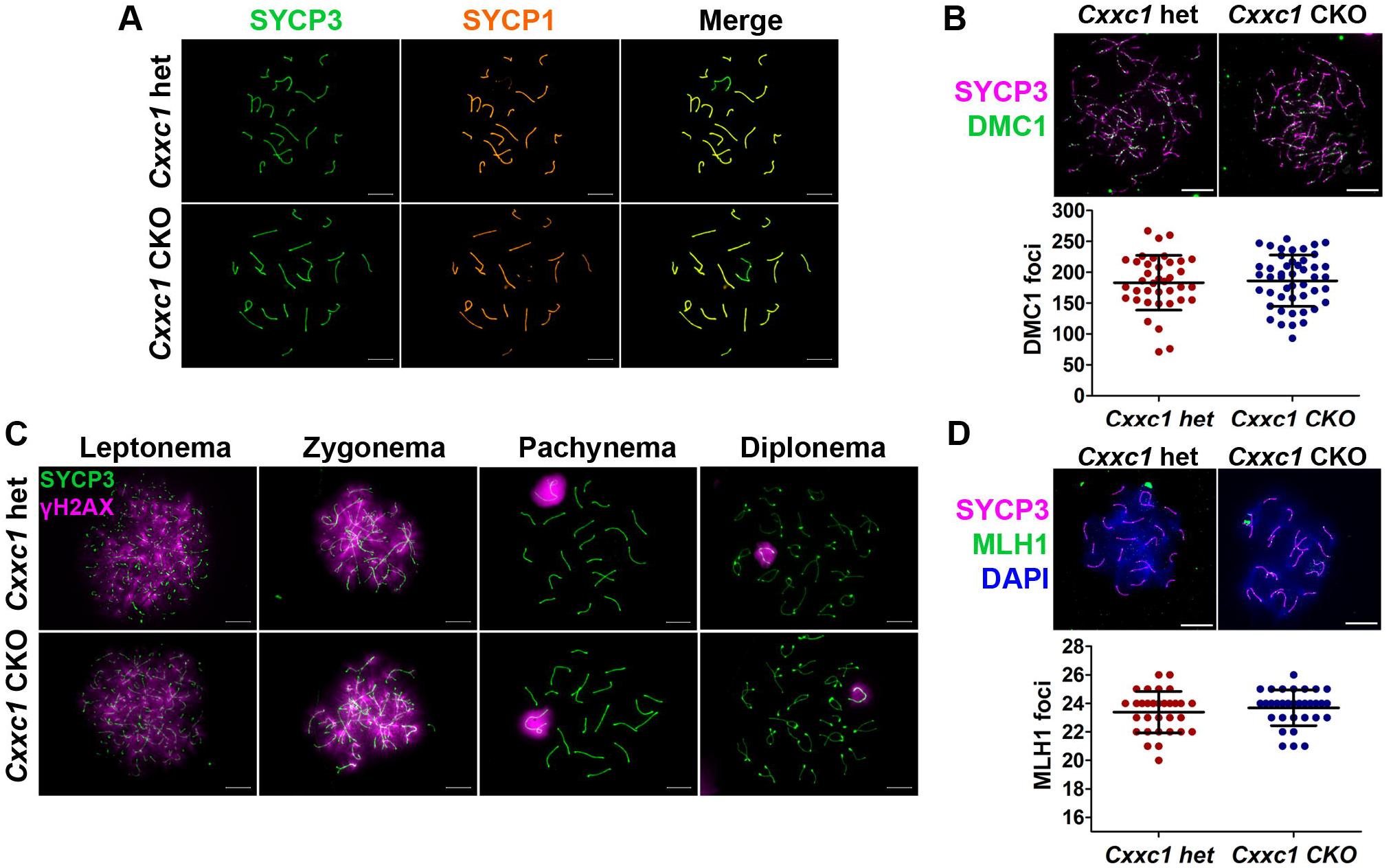
No major meiotic defects observed in *Cxxc1* CKO testis. (A) Immunostaining of SYCP3 and SYCP1 on adult *Cxxc1* het and CKO chromosome spreads. Green, SYCP3; orange, SYCP1. Scale bar, 10 μm. (B) DMC1 staining on *Cxxc1* control (n = 38) and CKO (n = 47) chromosome spread. Scale bar, 10 μm. Distribution plot of DMC1 foci in leptotene and zygotene spermatocytes was shown on the lower panel. Bars represent mean ± SD. *p* = 0.73 by Student’s t-test. (C) Immunostaining of SYCP3 and γH2AX on adult *Cxxc1* het and CKO chromosome spreads. Green, SYCP3; magenta, γH2AX. Scale bar, 10 μm. (D) Crossover number was measured by MLH1 staining on chromosome spreads of adult *Cxxc1* het (n = 31) and CKO (n = 32) spermatocytes. Magenta, SYCP3; green, MLH1; blue, DAPI. The distribution plot of MLH1 foci in pachytene spermatocytes was shown on the lower panel. Bars represent mean ± SD. *p* = 0.38 by Student’s t-test.

DSB formation was not affected by the loss of CXXC1 either, when measured by the number of foci of DMC1, the protein that binds to the single stranded DNA tails at DSB sites [7,8] (Fig 5B). We used staining for phosphorylated H2AX (γH2AX), which marks unrepaired DNA lesions, to test for the processing of recombination repair. The pattern of γH2AX staining was not changed in CKO compared to the het control spermatocytes, showing γH2AX signal throughout the nucleus in leptotene spermatocytes when DSBs occur, which was then restricted to the sex body in pachytene spermatocytes when the autosomal breaks are repaired (Fig 5C).

Finally, we examined whether loss of CXXC1 affects crossover determination process. Using MLH1 as a marker of crossover sites, we did not find any significant change of crossover number in the CKO spermatocytes compared to the het controls (Fig 5D).

Taken together, these results suggest that even though CXXC1 interacts with PRDM9 and H3K4me3 in spermatocytes, it is not required for PRDM9 binding at hotspots, their subsequent activation by PRDM9-dependent H3K4 trimethylation, DSB formation, repair, or crossover formation, and is therefore redundant for meiotic recombination events.

## Discussion

In this study, we demonstrate that CXXC1 interacts with PRDM9 in spermatocytes. However, CXXC1 is not a predominant interactor of PRDM9 and its binding to PRDM9 depends on the catalytic activity of PRDM9. The germ cell specific *Cxxc1* knockout male mice are fertile. In the knockout spermatocytes, the expression and function of PRDM9 are unchanged. The loss of CXXC1 does not affect DSB formation and repair, chromosome pairing and synapsis, and crossover numbers. Together, these results convincingly show that CXXC1 is redundant for normal meiotic recombination events and spermatogenesis.

The yeast CXXC1 ortholog Spp1 is a key player in recombination by linking H3K4me3 sites with the chromosome axis and connecting them with the recombination protein Mer2 [17–19]. These observations suggested that CXXC1 might play similar function in mammalian meiosis. However, our results show that CXXC1 is not an essential player in mammalian recombination where PRDM9 controls the initial recognition and activation of recombination hotspots, because in the absence of CXXC1, hotspot activation, axis integrity, DSB formation and crossover resolution occur normally. This suggests that PRDM9 dependent DSB formation and recombination determination pathway in most of mammals differs from that in the budding yeast. In species with functional PRDM9, the function of RMM complex consisting of orthologs of the yeast Rec114, Mei4 and Mer2 (REC114, MEI4 and IHO1 in mice) is still conserved [25,28–30], and the association between hotspots and chromosome axis is crucial for efficient DSB formation [25,31,32]. However, the interaction between CXXC1 and IHO1 does not seem to play the same functional role as the one between Spp1 and Mer2 in yeast. One important difference is that in organisms that do not use PRDM9, DSB occur at H3K4me3 sites, whereas in those that use PRDM9, DSB occur at hotspots where surrounding nucleosomes are methylated at both H3K4 and H3K36 [5,6]. This raises the likelihood that proteins with H3K36me3 methyl-reading activity, such as PWWP domain containing proteins [33], or with both H3K4me and H3K36me binding capability, such as Tudor domain containing proteins [34], might be involved in hotspot recognition in these species. Alternatively, activated hotspots may be recruited to the chromosome axis and DSB machinery without assistance of an H3K4me3/H3K36me3 reader. A recent study demonstrated that randomized DSBs induced by radiation in *Spo11* mutant spermatocytes are associated with chromosome axis and can successfully recruit DSB repair proteins such as DMC1/RAD51 complex [35]. Other direct PRDM9 interactors, such as EWSR1, EHMT2, and CDYL, could also be involved in hotspot association with the chromosome axis [15].

An alternative, PRDM9-independent pathway, can explain the fraction of DSB detected at promoters in wild type mice, and all DSB in PRDM9 mutant mice [8,36]. This pathway could involve CXXC1 as part of the SETD1 complex, known to bind H3K4me3 at promoters, in a way similar to Spp1-Mer2 role in yeast meiosis. It is not an essential pathway in most organisms using PRDM9 as hotspot determinant, but may play a major role in those lacking PRDM9, such as canids [37–39], where recombination hotspots are enriched in CpG-rich regions with a preference for unmethylated CpG islands [8,16,37,40], similar feature as CXXC1 binding sites [20,41]. A recent finding of a woman, who has no active PRDM9 but is fertile, suggests that this pathway may become activated and ensure proper recombination even in organisms using PRDM9 as a recombination regulator [42].

## Methods

### Mouse models

All wild-type mice used in this study were in the C57BL/6J (B6) background. Exon 2 and exon 3 of *Cxxc1* were flanked by two loxP sites using CRISPR. The *Cxxc1* conditional knockout mice used in this study were produced by a two-step deletion scheme. Mice that harbor two conditional *Cxxc1* alleles (*Cxxc1^loxp/loxp^*) were mated to Tg(Sox2-cre)1Amc/J mice (stock #004783) to generate one *Cxxc1* allele deleted mice. The *Cxxc1* hemizygous mice (*Cxxc1^Δ/+^*) were mated to Tg(Stra8-icre)1Reb/J (stock #08208) and Tg(Ddx4-cre)1Dcas/KnwJ (stock #018980) to obtain *Cxxc1^Δ/+;Stra8-iCre^* and *Cxxc1^Δ/+;Ddx4-Cre^* mice. Those *Ewsr1^Δ/+;Stra8-iCre^* and *Cxxc1^Δ/+;Ddx4-Cre^* mice were then mated to homozygous *Cxxc1* loxp mice to generate heterozygous control mice (*Cxxc1^loxp/+;Stra8-iCre^* and *Cxxc1^loxp/+;Ddx4-Cre^* designated as *Cxxc1* controls) or conditional knockout mice (*Cxxc1^loxp/Δ;Stra8-iCre^* and *Cxxc1^loxp/Δ;Ddx4-Cre^* designated as *Cxxc1* CKO).

B6(Cg)-Prdm9tm2.1Kpgn/Kpgn mice (*Prdm9^Set-/Set-^*) were generated in a previous study (unpublished data). B6;129P2-*Prdm9^tm1Ymat^*/J mice (*Prdm9^-/-^*) have been previously described [43]. All animal experiments were approved by the Animal Care and Use Committee of The Jackson Laboratory (Summary #04008).

### Co-immunoprecipitation assays

The co-IPs for PRDM9 and EWSR1 with testicular extract were carried out using our reported protocol [15]. Testicular total protein was extracted from twenty 14 dpp B6 and *Prdm9^Set-/Set-^* with 1 ml of Pierce IP buffer (Thermo Fisher Scientific, 87787). 10% of extract was set apart as input. The co-IP was perform by incubating extract with protein G Dynabeads conjugated with antibodies against PRDM9 (custom-made) or guinea pig IgG overnight at 4°C. Then, the beads were washed three times with 1 ml of Pierce IP buffer, eluted with 200 μl of GST buffer (0.2 M glycine, 0.1% SDS, 1% Tween 20, pH 2.2) for 20 min at room temperature and neutralized with 40 μl of 1 M Tris-HCl, pH 8. After heated at 95°C for 5 min, 10 μg of IP and input samples were then subjected to electrophoresis and western blotting for detection of PRDM9 (1:1000, custom made), EWSR1 (1:1000, Abcam, ab54708), CXXC1 (1:1000, Abcam, ab198977) and CTCF (1:1000, Abcam, ab70303).

The co-IP for CXXC1 was performed with isolated germ cells. Briefly, the seminiferous tubules were digested with liberase and the germ cells were isolated. Then, the nuclei were isolated by incubation germ cells in hypotonic lysis buffer (10 mM Tris-HCL pH 8.0, 1 mM KCl, 1.5 mM MgCl_2_) for 30 min at 4°C and spinning down at 10,000 *g* for 10 min. The nuclear extract was obtain by incubation with the nuclei lysis buffer (50 mM HEPES, pH 7.8, 3 mM MgCl_2_, 300 mM NaCl, 1 mM DTT and 0.1 mM PMSF), 5 U/μl DNaseI and 2 U/μl Turbonuclease for 30 min at 4°C. 10% of extract was saved as input. The co-IP was perform by incubating extract with protein G Dynabeads conjugated with antibodies against CXXC1 (Abcam, ab198977) or guinea pig IgG overnight at 4°C. After wash and elution, the IP and input samples were then subjected to electrophoresis and western blotting for detection of CXXC1 (1:1000, Abcam, ab198977), H3K4me3 (1:1000, Millipore, #07-473) and H3K9me3 (1:1000, Active Motif, 39766).

### Measurement of testis index

Testicular weight and body weight of adult B6 (n = 3), *Cxxc1* het (n = 3) and CKO (n = 4) mice were measured. Testis index was calculated as testis weight/body weight. Student’s t-test was used to determine the statistical significance.

### Fertility test

Fertility test was performed with 3 *Cxxc1* het control and 5 CKO male mice. Each mouse was mated with at least two B6 females for at least two month period. Litter size and viable pup number were record.

### Histology

Testis and epididymis from adult *Cxxc1* het control or CKO mice were dissected out, fixed with Bouin’s solution, and embedded in paraffin wax, and sectioned at 5 μm. Sections of testis were stained with Periodic acid–Schiff–diastase (PAS), and section of epididymis were stained with haematoxylin and eosin (HE) using standard techniques.

### Chromosome spread and FISH

The drying-down technique [44] was used for preparation of chromosome spreads from spermatocytes of 14-dpp and 8-weeks B6, *Cxxc1* control or CKO mice. Chromosome spread slides were immunolabeled with anti-PRDM9 (1:200), CXXC1 (1:1000), SYCP1 (1:300, Novus, NB300-229), SYCP3 (1:400, Novus, NB300-231), γH2AX (1:1000, Abcam, ab26350), DMC1 (1:200, Santa Cruz, sc-8973) or MLH1 (1:100, BD Pharmingen, 550838) antibodies.

### Immunofluorescence

For protein immunolocalization on tissue sections, testicular tissues from 8 week old B6, *Prdm9^-/-^, Cxxc1* control and CKO mice were dissected out, fixed with 4% paraformaldehyde solution overnight, embedded in paraffin wax. 5-μm sections were prepared. For antigen retrieval, sections were heated in a microwave in Tris-EDTA buffer (10mM Tris, 1mM EDTA and 0.05% Tween 20, pH 9.0) for 10 min and cooled down to room temperature. Then, sections were treated with PBS containing 0.1% Triton X-100, blocked with 10% normal donkey serum, and stained with antibodies against EWSR1 (1:200), PRDM9 (1:200), CXXC1 (1:1000).

### Chromatin immunoprecipitation and real-time PCR

Chromatin immunoprecipitation (ChIP) wad performed as previously described [45]. Briefly, spermatocytes were isolated from 14-dpp B6, *Cxxc1* het and CKO spermatocytes, and crosslinked using 1% formaldehyde. Nuclei were isolated using hypotonic lysis buffer (10 mM Tris-HCL pH 8.0, 1 mM KCl, 1.5 mM MgCl_2_) and digested by MNase. The ChIP was done using antibody against H3K4me3 (Millipore, #07-473). Real-time PCR was performed with purified ChIP DNA using Quantifast SYBR Green PCR Kit (Qiagen) Primer sequences used for realtime PCR are: *Pbx1*_F: AGAAACTGACATATGAAGGCTCA; *Pbx1*_R: GCTTTTGCTCCCTTAAACTGG; *Fcgr4*_F: CAAGGTGCATTCTTAGGAGAGA; *Fcgr4*_R: TTAATGCTTGCCTCACGTTC; *Hlx1*_F: GGTCGGTGTGAGTATTAGACG; *Hlx1*_R: GGCTACTATACCTTATGCTCTG; *Actinb*_promoter_F: GCCATAAAAGGCAACTTTCG; *Actinb*_promoter_R: TTTCAAAAGGAGGGGAGAGG; *Sycp3*_promoter_F: AAGGCGCCACAACCAAGG; *Sycp3*_promoter_F: TGCCTGGATGCCCAACTC.

## Acknowledgments

We thank all members of Petkov and Paigen labs, Mary Ann Handel and Ewelina Bolcun-Filas for their helpful comments, and Anita Adams for technical help.

## Author contributions

HT and PMP designed the experiments. HT and TB performed the experiments. HT and PMP analyzed the results. HT and PMP wrote the paper.

## Supporting Information

**Fig S1.**
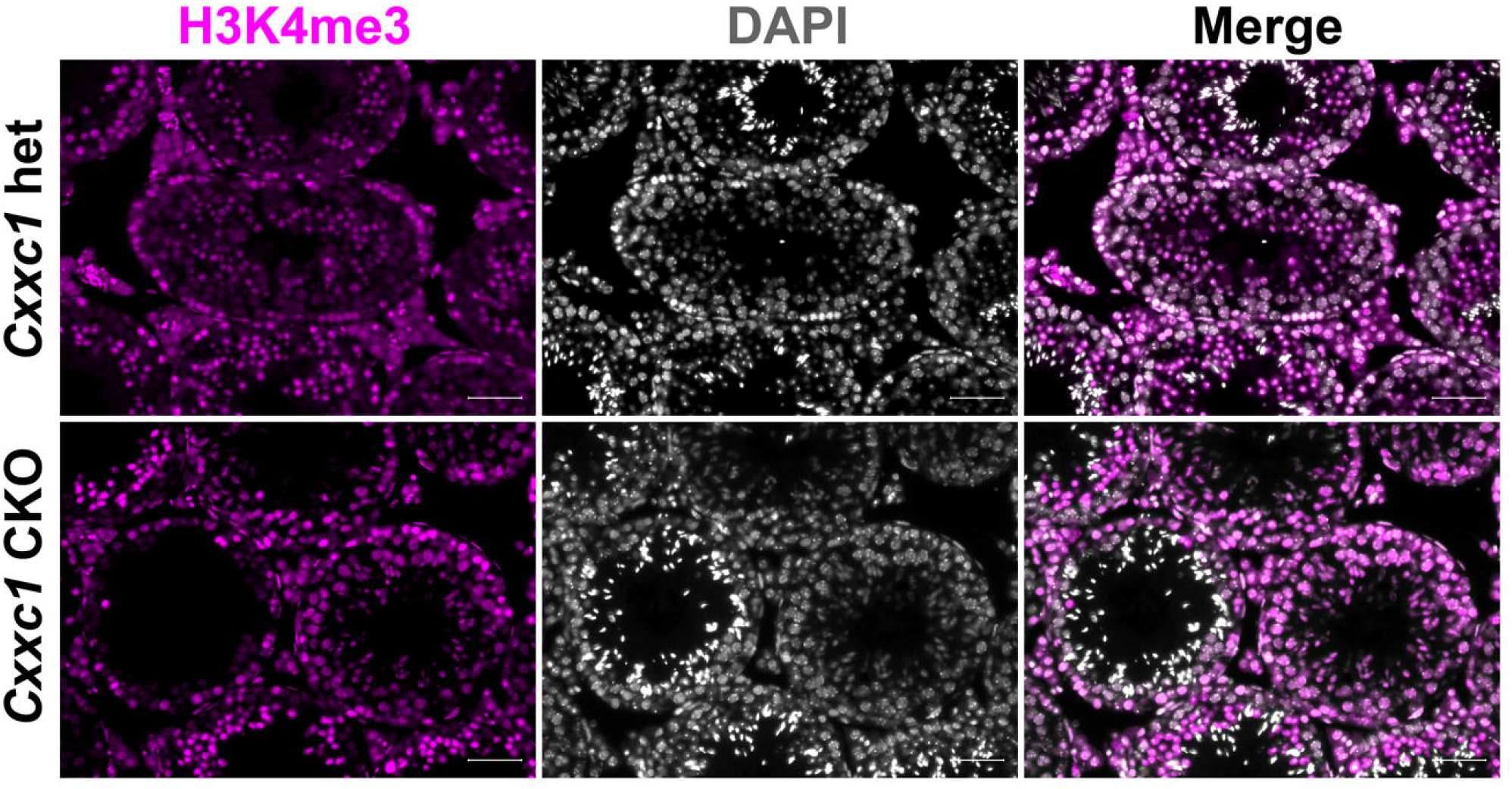
H3K4me3 staining on *Cxxc1* CKO seminiferous tubule cross sections. Immunofluorescence staining of H3K4me3 on adult *Cxxc1* het and CKO seminiferous tubule cross sections. Magenta, H3K4me3; Gray, DAPI. Scale bars: 50 μm.

